# Nanoscale imaging of expanded cells and proteins with spontaneously blinking dyes

**DOI:** 10.64898/2026.02.23.707413

**Authors:** Danush Taban, Marvin Jungblut, Made Budiarta, Dominic A. Helmerich, Cerridwen Kiesel, Sarah E. Plutkis, Luke D. Lavis, Donatus Krah, Ali H. Shaib, Sören Doose, Philip Kollmannsberger, Silvio O. Rizzoli, Gerti Beliu, Markus Sauer

**Affiliations:** Department of Biotechnology and Biophysics, Biocenter, University of Würzburg; Würzburg, Germany; Rudolf Virchow Center, Research Center for Integrative and Translational Bioimaging, University of Würzburg; Würzburg, Germany; Janelia Research Campus, Howard Hughes Medical Institute, Ashburn, VA, USA; Department of Neuro- and Sensory Physiology, University Medical Center Göttingen, Göttingen, Germany; Faculty of Chemistry and Pharmacy, Regensburg Center for Ultrafast Nanoscopy (RUN), Universitätsstrasse 31, 93053 Regensburg

## Abstract

Expansion microscopy (ExM) enables nanoscale imaging on standard microscopes, but combining ExM with single-molecule localization microscopy (SMLM) remains difficult, owing to the incompatibility of expanded hydrogels with photoswitching buffers. Here, we introduce a single-step expansion microscopy method that allows SMLM with spontaneously blinking dyes in 6-14× expanded samples, without re-embedding. We demonstrate nanometer-resolution imaging by resolving the organization of the nuclear pore complex (NPC) and the molecular structure of recombinant homotrimeric proliferating cell nuclear antigen (PCNA).

## Main text

Single-molecule localization microscopy (SMLM) provides nanometer-scale spatial resolution but typically requires fluorophores that undergo photoswitching or photoactivation in special buffers^1-3^. Alternatively, transient binding approaches such as DNA-PAINT with imager strands can be used but also requires aqueous buffers with high salt concentrations^4^. Recently the resolution limits of SMLM have been further improved demonstrating localization precisions in the Ångström range on isolated molecules^5,6^. In parallel, expansion microscopy (ExM) emerged as an alternative super-resolution microscopy method^7^. By physically expanding biological specimens after embedding in a swellable hydrogel, ExM enables super-resolution imaging on standard microscopes without the need for specialized optics or sophisticated imaging conditions. In addition, post-expansion labeling methods have been developed, allowing improved epitope accessibility and thus higher label densities, while simultaneously reducing the linkage error^8-10^. Protocols have been improved and expansion factors continuously increased promising theoretically unlimited spatial resolutions^11-15^. However, higher expansion factors often come at the cost of technical complexity, particularly concerning the handling and advanced imaging of large hydrogels.

Alternatively, imaging of moderately expanded and thus manageable hydrogels by SMLM can push the spatial resolution towards the molecular resolution range. Using expanded samples for imaging the distance between adjacent fluorophores is large enough to prevent unintended interfluorophore energy transfer that results in accelerated photobleaching and lower localization probability^16^. However, performing *d*STORM in expanded samples requires photoswitching buffers; this can be accomplished by re-embedding in an uncharged hydrogel, which maintains mechanical stability and avoids shrinking of the expanded gel^17^. In addition, the most popular *d*STORM dyes Cy5 and Alexa Fluor 647 are efficiently destroyed during re-embedding^18^. Therefore, previous implementations of Ex-*d*STORM achieved expansion factors of 3-4-fold and provided only modest resolution improvements^17,19-21^. Recently, one-step nanoscale expansion microscopy (ONE) demonstrated imaging of the shape and conformation of single recombinant and isolated proteins at nanometer resolution^22^. ONE uses fluorescence intensity fluctuations of dyes in water in ∼10-fold expanded samples. After anchoring and proteinase K digestion the newly created amino groups are labeled with NHS-functionalized fluorophores^23^. Although ONE avoids re-embedding of the expanded sample in a neutral gel, the fluorescence fluctuations of fluorophores is this method cannot be used for SMLM.

Here, we introduce a simplified and fast single-step TREx-based expansion protocol^13^ that provides ∼6× expansion of cellular multiprotein complexes combined with ultrastructure preservation, improved accessibility and integrity of epitopes, and ∼14× expansion of isolated proteins. Using JF_635_b, a spontaneously blinking hydroxymethyl-Si-rhodamine (HMSiR) analog based on a bright Janelia Fluor (JF) dye^24^, re-embedding can be avoided allowing us to resolve the molecular shape of isolated JF_635_b-NHS labeled proteins and image nuclear pore complexes (NPC) in intact cells with molecular resolution using a simplified workflow that can be completed within two days. This protocol allows Ex-SMLM in water without dedicated photoswitching buffers and converts standard widefield illumination into a robust single-step route to molecular resolution (Fig. 1a).

**Figure 1.**
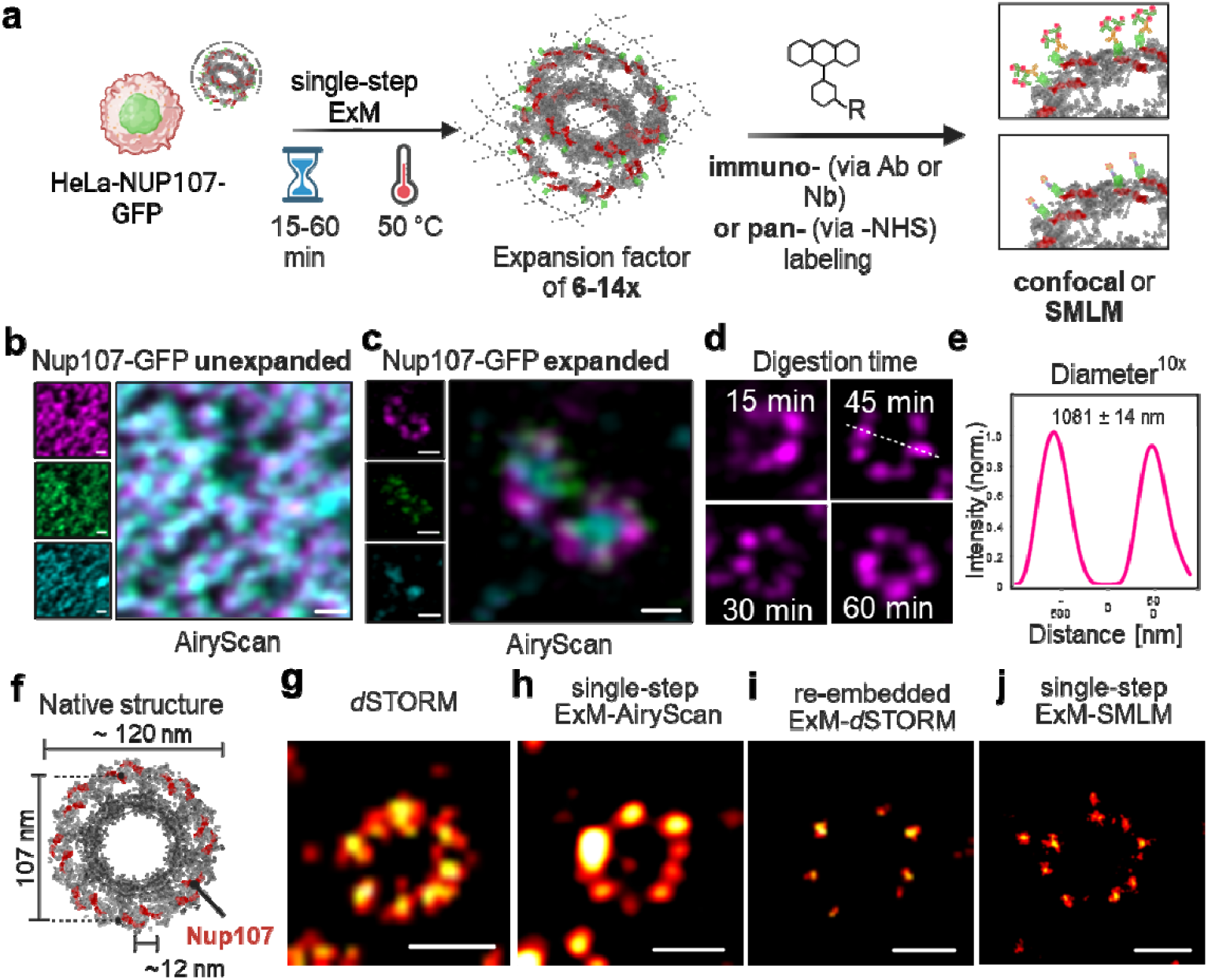
Single-step Ex-SMLM enables rapid expansion with preserved ultrastructure. **a**, Schematic of the single-step expansion workflow. Cellular protein complexes are anchored, polymerized, and digested for 15–60 min at 50 °C, yielding expansion factors of ∼6–14 depending on sample type. Expanded hydrogels are compatible with NHS labeling and post-expansion immunolabeling and can be imaged directly in water using confocal microscopy or SMLM. **b**, Pre-expansion Airyscan image of HeLa cells expressing Nup107– GFP, labeled with biotinylated anti-GFP nanobody and ATTO643-streptavidin. The inset shows individual channels for GFP (green), WGA (cyan), nanobody (magenta), and merged view. **c**, Post-expansion Airyscan image of NUP107-GFP NPCs labeled after expansion, demonstrating preserved GFP fluorescence and epitope integrity. The inset displays the individual fluorescence channels as shown in (**b**). **d**, Airyscan images of individual NPCs after single-step expansion with digestion times of 15, 30, 45, or 60 min at 50 °C. Even shorter digestion times of 15–30 min preserve the ultrastructure and allow full hydrogel expansion. **e**, Average radial intensity profile of expanded NPC rings after 45 min digestion showing a peak-to-peak diameter of 1081 ± 14 nm (mean ± s.d.) in n = 12 cells, corresponding to an effective expansion factor of approximately 10-11 for pre-expansion labeling. **f**, NPC structure highlighting the organization of NUP107 subunits (red), including the ∼12 nm distance between adjacent monomers in NUP107 dimers and the ∼107 nm diameter of the NPC ring. **g–j**, Comparison of NPC imaging methods. **g**, *dS*TORM of unexpanded NPCs labeled with AF647 anti-GFP nanobody resolves the characteristic eightfold symmetry. **h**, Airyscan image of a single-step expanded NPC reveals well-defined NPC rings at ∼10x expansion. **i**, Ex-*d*STORM image of a re-embedded ∼5.3× expanded NPC. **j**, Single-step Ex-SMLM image measured in pure water with spontaneously blinking dye JF_635_b visualizes the eightfold NPC symmetry at ∼6× expansion without re-embedding or addition of switching buffer. Scale bars: 500 nm (**b**), 500 nm (**c**), 100 nm (**g**), 1000 nm (**h**), 400 nm (**i**) and 300 nm (**j**).

To validate ultrastructure preservation, we used the NPC as a well-established reference structure. NPCs are among the largest proteinaceous assemblies in the cell and are constructed of around 30 different proteins called nucleoporins (NUPs) that assemble in multiple copies to form a complex with eight-fold rotational symmetry^25^. For the human NPC, electron microscopy^26^ and cryo-electron microscopy^27^ have provided a high-resolution structural map of the structure and most of the nucleoporin components. Here, we used a previously described HeLa cell line with stable expression of NUP107-GFP to investigate the expansion factor, structural distortions and labeling efficiency^28^. NUP107 is a nucleoporin present in 32 copies per NPC. It forms a cytoplasmic and nucleoplasmic ring, consisting of 16 NUP107 copies, respectively. Each of the two rings contains 8 NUP107 dimers spaced 12 nm apart^26^. As prolonged digestion can compromise ultrastructure, we first systematically optimized the proteolysis step. Shortening the TREx digestion time to 30 min at 50 °C preserved GFP fluorescence and NPC symmetry while still allowing maximal hydrogel expansion (Fig. 1b-d and Supplementary Fig. 1). Notably, extending the digestion to overnight conditions caused substantial disruption of NPC ring geometry and a pronounced reduction in both GFP and streptavidin signal, underscoring the need for controlled digestion times (Supplementary Fig. 2). Pre-expansion labeling with biotinylated anti-GFP nanobody and ATTO643-streptavidin, followed by TREx embedding, yielded peak-to-peak NUP107 distances of ∼1.0 µm corresponding to an effective expansion factor of 10 to 11 for NPCs (Fig. 1e and Supplementary Fig. 1).

**Figure 2.**
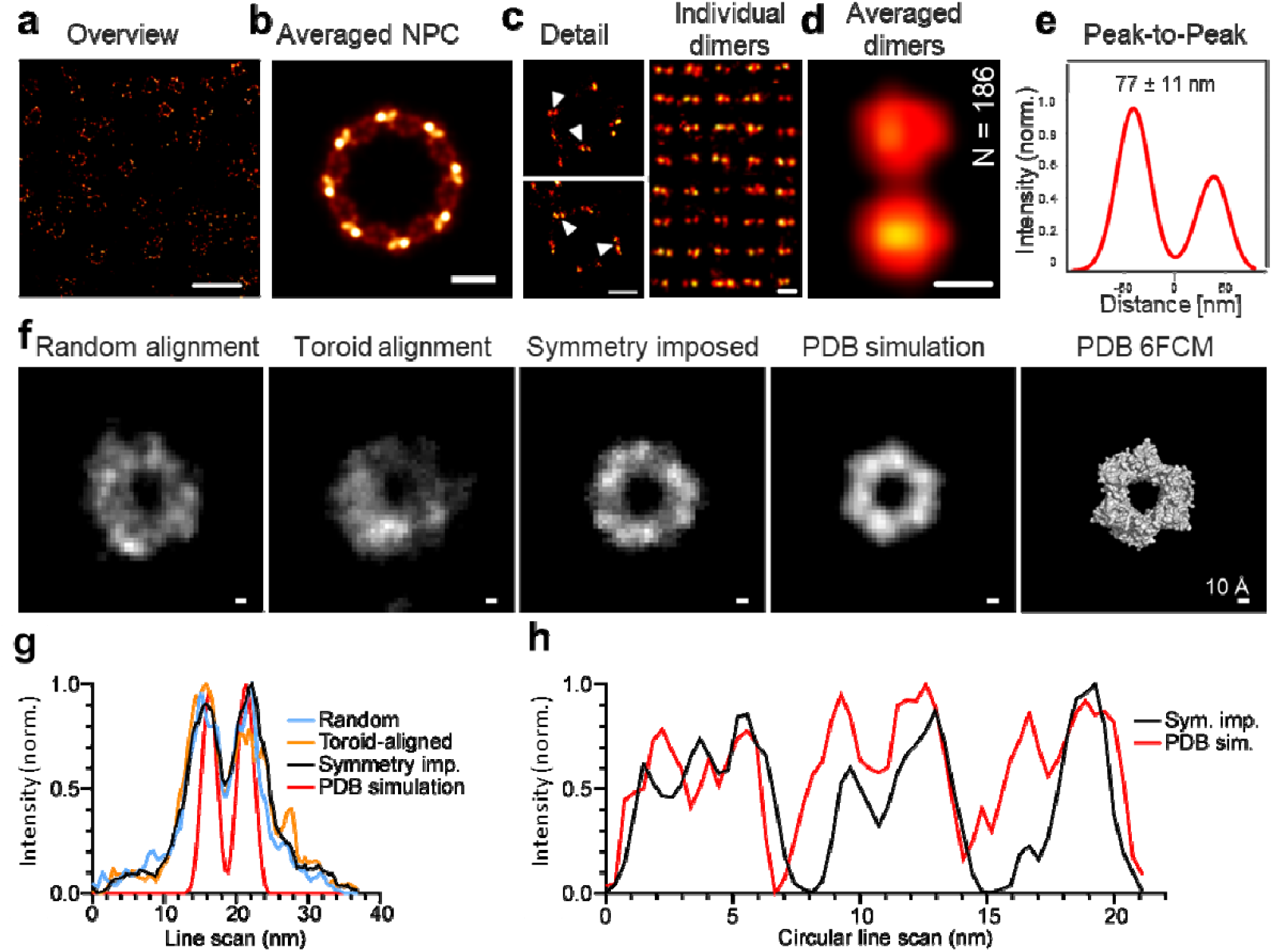
Single-step Ex-SMLM resolves the molecular architecture of NPCs and isolated PCNA at nanometer resolution. **a**, Single-step Ex-SMLM image of Nup107-labeled NPCs in expanded HeLa cells, showing high labeling density and uniform ring morphology across the field of view. **b**, Particle-averaged NPC reconstruction from 183 individual complexes, revealing the characteristic eightfold symmetry and preserved ring geometry. **c**, Zoom-ins of individual NPCs (arrowheads) showing clearly separated Nup107 dimer signals. **d**, Gallery of individual resolved Nup107 dimers (left) and the corresponding average dimer structure (right; n = 186). **e**, Radial intensity profile of the average dimer pair showing a peak-to-peak distance of 77 ± 11 nm (s.d.), corresponding to a native spacing of ∼12 nm at ∼6.4× microscopic expansion. **f**, Images of PCNA proteins labeled with JF_635_b-NHS, after expansion (Supplementary Fig. 5) were centered, overlaid and summed (leftmost panel, random alignment, n = 81). The images were subjected to alignment to a toroid structure, followed by 3-fold symmetry imposition. For comparison, the expected fluorescence pattern provided by the known coordinates of PCNA (PDB structure 6FCM)^31^ was simulated, and a surface view of the structure is also shown (rightmost panel). **g**, Horizontal line scans performed across the images from panel f. **h**, Radial intensity profiles for the PCNA reconstruction with imposed symmetry and for the PDB-simulated PCNA image. Similar patterns are observed (Pearson’s correlation coefficient of 0.60, p = 0.000000604). Scale bars: 2 µm (**a**), 200 nm (**b**) 300 nm (**c**), 100 nm for individual and 50 nm for average dimers (**d**), and 1 nm (10 Å) (**f**).

Using CF568-labeled wheat germ agglutinin WGA-CF568 to label the central channel of NPCs, airyscan microscopy easily resolved the nucleoplasmic and cytoplasmic ring of the NPC with WGA being localized in between the inner and outer layer (Extended Data Fig. 1).

To quantify the performance Ex-SMLM with JF_635_b, we first performed reference *d*STORM experiments on unexpanded NPCs. We labeled unexpanded HeLa-NUP107-GFP cells with an AF647 labeled anti-GFP nanobody (anti-GFP nanobody-X4-AF647). *d*STORM images revealed the 8-fold symmetry of the NPC and exhibited an average peak-to-peak distance of 105.8 ± 6.8 nm (mean ± s.d.), which corresponds well to the expected distance of 107 nm^26^ (Fig. 1g and Supplementary Fig. 3). Re-embedding expanded samples in a neutral gel enabled Ex-*d*STORM of NPCs with AF647 but reduced the effective expansion factor to around 5.3 (Fig. 1i and Supplementary Fig. 4), highlighting the practical limitations of buffer-based *d*STORM on expanded hydrogels.

We then tested whether post-expansion labeling with a spontaneously blinking dye can be used advantageously to achieve molecular resolution using a single TREx expansion step. Post-expansion immunostaining with rabbit anti-GFP and JF_635_b-labeled secondary antibodies followed by SMLM in water produced robust blinking and high localization densities across expanded NPCs (Fig. 1j, Extended Data Fig. 2 and Supplementary Movie S1). The NUP107-JF_635_b signals showed a mean peak-to-peak-distance of 663 nm ± 79.6 nm (s.d.) resulting in an average expansion factor of 6.2× with a clear preservation of the NUP107 subunits (Figs. 2a,b and Extended Data Fig. 2). Furthermore, Ex-SMLM allowed us to clearly resolve NUP107 dimers separated by only 12 nm (Fig. 2c,d). The average measured distance of 77±10.7 nm (mean ± s.d.) corresponds to a microscopic expansion factor of 6.4× (Fig. 2e). Particle averaging of 183 NPCs further confirmed the eightfold symmetry of dimers and revealed consistent substructure within NUP107 clusters, demonstrating that single-step Ex-SMLM with JF_635_b enables molecular resolution imaging of multiprotein complexes in cells.

To test whether the same workflow can retrieve the molecular shape of isolated proteins, we turned to proliferating cell nuclear antigen (PCNA), which has previously been used as a protein-based optical ruler^29^. PCNA plays an important role in DNA replication and repair. It forms a homotrimeric ring with inner and outer diameters of 3.4 nm and 8.0 nm, respectively^30^. We anchored purified PCNA to the TREx gel using Acryloyl-X and, after digestion, labeled newly exposed peptide termini with JF_635_b-NHS. Ex-SMLM of maximally expanded gels revealed ring-like structures with a central hole (Fig. 2f-h, Supplementary Fig. 5 and Supplementary Movie S2). From the averaged image we determined an expansion factor of 13.6× for isolated PCNA proteins which is slightly higher than the expansion factors achieved for cellular proteins (NPCs) using the same protocol. This demonstrates that, for isolated proteins, single-step Ex-SMLM can approach the structural information content typically associated with electron microscopy, while remaining compatible with standard fluorescence microscopes.

To summarize, we developed a modified ExM protocol based on a simplified version of TREx^13^, which substantially minimizes the time and complexity typically associated with high-resolution ExM workflows. Our approach combines shorter digestion times and anchoring steps with the use of JF_635_b, a spontaneously blinking fluorophore for SMLM^24^, altogether enabling sub-10 nm resolution fluorescence imaging in cells within a two-day experimental timeline.

In practice, the method condenses ExM sample preparation to a single expansion step, removes the need for re-embedding and specialized blinking buffers, and still preserves the ultrastructure and epitope integrity of cellular and purified protein targets. In addition to NPCs and isolated proteins, we also assessed whether clathrin-coated pits (CCPs) retain their characteristic ultrastructure after single-step expansion. Expanded CCPs immunolabeled post-expansion revealed clear triskelion organization (Extended Data Fig. 3), underscoring that the protocol preserves structurally complex endocytic assemblies. The higher microscopic expansion of isolated PCNA compared to NPCs suggests that intermolecular constraints in crowded cellular environments currently limit the achievable expansion factor for large multiprotein complexes but at the same time underline the potential of improved hydrogel formulations and digestion strategies.

Together, these results establish Ex-SMLM with JF_635_b as a simple and robust route to single-step expansion-based SMLM that resolves multiprotein complexes in cells with molecular resolution and structures of isolated multimeric protein. We anticipate that systematic optimization of spontaneously blinking dyes and hydrogel chemistries will further broaden the scope of this approach and enable routine molecular structure imaging in cells.

## Acknowledgements

The authors thank Elke Maier and Ivan Simeonov for cell culture support. D.T., M.J., D.A.H., C.K., S.D., S.O.R., A.H.S., D.K. and M.S. acknowledge funding from the European Research Council (ERC) under the European Union’s Horizon 2020 research and innovation programme (grant agreement No 835102). S.E.P and L.D.L. are funded by the Howard Hughes Medical Institute (HHMI). This project is supported by the Federal Ministry for Economic Affairs and Climate Action (BMWK) based on a decision by the German Bundestag (Grant Agreement No. KK5665801HV4 to G.B.).

## Author contributions

D.T., G.B., and M.S. conceived and designed the project. S.D., G.B. and M.S. supervised the project. D.T., C.K. and M.J. prepared samples. D.T., M.J. and D.A.H. performed all *d*STORM experiments. S.E.P. and L.D.L. provided spontaneously blinking dyes. A.H.S., S.O.R., D.K., P.K. and D.T. performed data analysis. M.B. provided purified PCNA. D.T., G.B., and MS. wrote the manuscript with input from all authors. All authors reviewed and approved the manuscript.

## Online Methods

### Cell culture

COS- 7 monkey kidney cells were purchased from CLS Cell Line Service GmbH and cultured at 37°C and 5% CO_2_ in DMEM with L-glutamine containing 10% FBS and 100 U/ml penicillin and 0.1 mg/ml streptomycin. 40,000 to 50,000 cells per well were seeded on round 12 mm high precision cover glasses (No. 1.5, Marienfeld, 0117520) in a 4-well culture plate (Techno Plastic Products, 92012) and grown 24 h prior to fixation. HeLa-NUP107-GFP-cells^28^ were cultured as described above. 150,000 cells per well were seeded on round 12 mm high precision cover glasses (No. 1.5, Marienfeld, 0117520) in a 4-well culture plate (Techno Plastic Products, 92012) and grown 24 h prior to fixation.

### Sample preparation. COS7-cells

To fix the cells, all solutions were pre-warmed to 37°C and fixation was carried out on a heating plate set to 37°C. Cells were fixed in cytoskeleton buffer (CB-buffer, 10 mM MES, 150 mM NaCl_2_, 5 mM EGTA, 5 mM glucose and 5 mM MgCl_2_, pH 6.1). Cells were fixed and permeabilized simultaneously. First incubating for 90 s in a primary fixative solution of 0.3% glutaraldehyde and 0.25% Triton X-100 (Surfact-Amps Detergent Solution (10% (w/v), 28313, ThermoFisher)) in CB-buffer followed by a second fixation step using 2% glutaraldehyde in CB-buffer for 10 min. The sample was then washed three times with PBS (1x) for 5 min each.

### Sample preparation. HeLa-NUP107-GFP

Fixation solution was pre-warmed to 37°C and fixation was carried out on a heating plate set to 37°C. Cells were fixed in 2.4% formaldehyde in CB-buffer for 30 min followed by a blocking and permeabilizing step using 5% bovine serum albumin (BSA, A2153, Sigma) in blocking solution (4 mM EDTA, 0,02% Tween-20 (10%, 93774, Sigma), 0.05% NaN_3_) in PBS (1×). The sample was then washed three times with PBS (1×) for 5 min each. Unless stated otherwise, all incubations were performed at room temperature. Immunostaining was carried out before gelation (prelabeling) or after gelation (postlabeling).

### ExM protocol

To copolymerize amine groups into the hydrogel, cells were incubated in the amine reactive agent Acryloyl-X, SE (6-((acryloyl)amino)hexanoic acid, succinimidyl ester (A20770)) (0.1 mg/ml) in 200 mM NaHCO_3_ for 2 h at room temperature. The reagent was prepared freshly from desiccated stock aliquots kept at −20 °C. Gelation was performed in a four-well plastic plate with a lid (Techno Plastic Products, 92012), serving as a humidified chamber. A layer of parafilm is placed in the middle two wells to provide a smooth and hydrophobic surface for gelation. Wet paper towels are placed in the outer two wells to provide humidity. 12 mm cover glasses with cells facing down were placed on a 50 µl pre-chilled ExM monomer solution (1.1 M sodium acrylate (SA, 97–99%, 408220, Sigma), 2.0 M acrylamide (AA, 40%, A4058, Sigma), 70 ppm *N,N*′-methylenebisacrylamide (BIS, 2%, M1533, Sigma)) supplemented with 0.15% *N,N,N*′,*N*′-tetramethylethylenediamine (TEMED, T7024, Sigma) and 0.15% ammonium persulfate (APS, A3678, Sigma). Samples were left on ice for 5 min before carefully transferring them to a humidified chamber and incubate for 1 h at 37 °C. After gelation samples were treated with 8 U/ml Proteinase K (P4850, Sigma) in improved digestion buffer^12^ (50 mM TRIS (T6066, Sigma), 800 mM guanidine HCl (50933, Sigma), 2 mM CaCl_2_, (Merck, 223506) and 0.5% (vol/vol) Triton X-100 (Thermo, 20770) in ddH_2_O with the pH adjusted to 8.1) depending on the target for 15 min–60 min at 50°C. After digestion, gels were washed with PBS (1x) three times for 5 min each before labelling either with antibodies, nanobodies or NHS-ester) initializing final expansion in double deionized water. Water was exchanged several times until the maximum expansion factor of the hydrogel was reached. Maximum expansion factor was reached when the gel-size did not change withing three water changes.

### Prestaining with WGA-CF568

After fixation, blocking and permeabilization, cells were incubated with 7.5 µg/ml Wheat Germ Agglutinin CF568 (WGA-CF568, 19077-1, Biotium) in 5% BSA in PBS (1×) for 2 h at room temperature, under gentle agitation. Cells were washed three times with PBS (1×). All prelabeled samples were postfixed with 2,4% formaldehyde for 15 min at room temperature, followed by crosslinking and gelation as stated above.

### Nanobody staining of unexpanded HeLa-NUP107-GFP cells

After fixation, blocking and permeabilization, cells were incubated in a staining solution, containing 100 nM anti-GFP nanobody X4-AF647 (N0304-AF647, NanoTag) in 5% BSA in PBS (1×) for 3h at room temperature or overnight at 4 °C. Samples were washed three times in PBS (1×) for 5 min each.

### Nanobody staining of expanded HeLa-NUP107-GFP cells

Following the ExM protocol, the sample was incubated in an nanobody staining solution, containing 500 nM anti-GFP nanobody X4-AF647 (N0304-AF647, NanoTag) or 500 nM anti-GFP nanobody X4-ATTO643 (N0304-At643) or 500 nM single domain antibody (sdAB) anti-GFP nanobody conjugated to biotin (N0305-BIOTIN, NanoTag) for 3h at room temperature or overnight at 4 °C under gentle agitation. Samples stained with sdAB anti-GFP nanobody conjugated to biotin were further poststained with Streptavidin (Invitrogen, S21374) conjugated to AF647 (20 µg/ml) over night at 4 °C under gentle agitation. Afterwards samples were washed three times in ddH_2_O with 0.2% Tween-20.

### Antibody staining of expanded HeLa-NUP107-GFP cells

Following the ExM protocol, the sample was incubated in an antibody staining solution, containing a primary rabbit-anti-GFP-antibody (1/250; Invitrogen, G10362) and secondary anti-rabbit-antibody (20 µg/ml) conjugated to ATTO643 of AF647 for 3 h at room temperature or overnight at 4 °C under gentle agitation. Samples were then washed three times in ddH_2_O with 0.2% Tween-20.

### Reembedding of expanded samples

Hydrogels are prone to shrinking if the osmolarity of the surrounding media differs from its own. To avoid shrinking and thus drift during measurements, caused by *d*STORM photoswitching-buffer, an uncharged gel was crosslinked throughout the hydrogel. It was chemically bound on a Silanized coverglass. Round 24 mm high precision coverglasses (No. 1.5, Marienfeld, 011264) were rinsed with ddH_2_O before sonicating in absolute ethanol for 10 min. After another rinse with ddH_2_O, the coverglasses got sonicated in 1 M potassium hydroxide (Merck, 105032) for 10 min before another rinsing with ddH_2_O and drying at 70 °C until dry. 10 ml of a working solution containing 10 µl (3**-**aminopropyl)triethoxysilan (APTES, 440140, Sigma), 200 µl glacial acetic acid (Sigma, A6283), 1,8 ml ddH_2_O in 8 ml absolute ethanol was distributed evenly on the cleaned 24 mm cover glasses and incubated for 1,5 h or until fully evaporated. Cover glasses were then rinsed with ddH_2_O and air-dried. After silanization, cover glasses were stored at −20 °C. To re-embed the expanded hydrogels, they were placed in a 6-well cell culture plate and each sample was covered with 3 ml of freshly prepared re-embedding solution (10% acrylamide, 0,15% *bis*-acrylamide, 0,03% TEMED, 0,03% APS in 5 mM Tris (pH 8.9)). The samples were incubated twice with freshly prepared solution for each 30 min under gentle agitation, to bring oxygen into the solution, to prevent it from polymerizing prematurely. After that, samples were placed on silanized coverglasses and transferred to a humidified container, equipped with gas injection holes. Oxygen was then removed by purging the chamber with nitrogen for 15 min. The samples were then incubated for 2 h at 37 °C. After the re-embedding gel was polymerized, samples were washed twice with ddH_2_O and placed in imaging buffer.

### SMLM of JF_635_b labeled samples

Super-resolution imaging of JF_635_b labeled samples was performed on a wide-field fluorescence microscope (IX-71, Olympus) using a 641 nm diode laser (Cube 640-100 C, Coherent), combined with a clean-up filter (Laser Clean-up filter 640/10, Chroma). The laser beam was focused onto the back focal plane of a 60x Oil-immersion objective (NA 1.45, Olympus) and the emission light was separated from the excitation light using a dichroic mirror (HC 560/659, Semrock). Further spectral filtering was done using a bandpass filter (FF01-679/41-25, Semrock). Images were captured using an electron-multiplying CCCD camera (iXon DU-897, Andor). For data analysis, the pixel size was determined to be 128 nm. *d*STORM measurements were recorded with 15.000 to 30.000 frames and an exposure time of 20 or 50 ms (corresponding to a frame rate of 50 or 20 Hz). Illumination intensity during measurement was approximately 0.75 kW/ cm^2^. Samples were imaged using a highly inclined and laminated optical sheet (HILO). SMLM measurements were performed in ddH_2_O and fully expanded hydrogels. All SMLM results were analyzed and reconstructed using rapidSTORM3.3 ^32^ and the localization precision was calculated according to Mortensen *et al*.^33^. In NUP107 Ex-SMLM and PCNA experiments we achieved lateral localization precisions of 12.9 ± 6.3 nm (mean ± s.d.) and 14.3 ± 7.5 nm (mean ± s.d.), respectively.

### Segmentation and averaging of NPCs

We performed automated segmentation and filtering of JF_635_b-labelled NPCs in six different fields of view (FOV) as described in Löschberger *et al*.^34^ and Thevatasan *et al*.^28^. To detect NPC candidates, a template image (Gaussian-blurred ring with 53nm radius) was convolved with a histogram image of the FOV, and local maxima spaced at least 1.5 NPC radii apart were selected as candidates. We then fit a circle to all localizations within a window of 150 nm × 150 nm around each detected NPC center and applied a set of filtering criteria following a similar approach as described in Thevatasan *et al*.^28^ to remove false positives: the circle radius must be within 50%-150% of 53 nm, each NPC must contain at least 20 and at most 5000 localizations, no more than 30% of localizations must be outside 64 nm and no more than 20% of localizations inside 42 nm, and at least 6 of 8 sectors of the circle must contain at least five localizations (at least 6 subunits should be labelled). After filtering, the NPCs were denoised using DBSCAN (eps = 30, min_samples = 3). An average NPC was calculated using template-free all-to-all particle fusion of n = 183 NPCs as described in Heydarian *et al*.^35^.

### PCNA Labelling

PCNA was expressed in C41(DE3) cells (Sigma Aldrich) and purified as previously described ^7^. To prepare PCNA for imaging, 100 µg of purified PCNA was crosslinked overnight with 0.3 mg/ml Acryloyl X-SE (Thermo, 20770) solution in 200 mM NaHCO_3_ in PBS (1×) at room temperature. Crosslinking was performed on cleaned 12 mm coverslips. After incubation, the crosslinking solution was left to evaporate under a fume hood until only a small residual volume remained, ensuring that the sample did not dry out completely. 60 µl of the already prepared TREx monomer solution (1.1 M (w/w) sodium acrylate (SA, #408220, Sigma), 2.0 M (w/w) acrylamide (AA, #A4058, Sigma), 50 ppm (w/w) *N,N’*-methylenebisacrylamide (bis), dissolved in 1× PBS. 0.15% TEMED and 0.15% APS) was then pipetted on parafilm and the PCNA coated coverslip was then flipped on it. Polymerization was allowed to proceed in a humidified chamber for 5 min on ice and then 1h at room temperature. For the homogenization of proteins and single molecules, samples were treated with 8 U/ml Proteinase K in improved digestion overnight at 50°C. After digestion, gels were washed with PBS (1×) three times for 5 min each before staining with 30 µg/ml JF_635_b-NHS in 200 mM NaHCO_3_ (pH. 8.1) for 3 h at room temperature before washing washed three times in ddH_2_O with 0.2% Tween-20.

PCNA was analyzed as follows. Regions of interest (ROIs) containing evident particles were identified by an automatic procedure, which combined a bandpass filter, to remove broad background signals, followed by a median filter, to remove noise pixels, and by a morphological erosion procedure, to remove objects substantially smaller than the expected size of PCNA (smaller than 8.0 nm at their widest point). The ROIs were then manually overlaid and centered, and they were summed, to provide an initial view of PCNA objects (Fig. 2a, random alignment). Alternatively, the ROIs were aligned to a toroid shape, with or without imposing a 3-fold symmetry, as previously described^36^. To compare the resulting images to the known PCNA structure, fluorophores were imposed on the coordinates of its amino acids, relying on PDB structure 6FCM as described before^22^. A noise level of up to 2 nm was enabled for this imposition, to account for imaging and/or ExM errors.

### Poststaining of nuclei with DAPI

Staining of the nucleus can still be performed after gelation and digestion of the hydrogels. To do this the gels, or sections of the gels are transferred into a new plastic dish or reaction tube and submerged in 1× PBS. DAPI in a 1 mg/ml stock solution in DMSO is added 1:500 to the submerged gels and incubated at room temperature under gentle agitation for 1 h. After staining the gels are washed 3–5 times in 1× PBS to wash out unbound DAPI.

### Equipment for AiryScan imaging of expanded gels

Expanded gels were cut to size and seeded on APTES (Sigma, A36480) coated 24 mm round high-precision coverslips (Marienfeld, 0117520). For imaging, these coverslips were transferred into 24 mm coverglass imaging chambers (Marienfeld, 0117640) Imaging was performed using a Zeiss LSM 900 with Airyscan 2 in super-resolution (SR) mode using a 63×/1.4 NA oil immersion objective for imaging of unexpanded samples, and a 40×/1.2 NA water immersion objective for imaging of expanded samples. For excitation 4 laser lines, 405 nm (30 mW), 488 nm (30 mW) and 639 nm (25 mW) diode lasers and a 561 nm (25 mW) DPSS laser are present. For each fluorophore in the sample appropriate laser lines and filter settings were applied, following the suggested optimal imaging settings provided by the microscopes control software ZEN 2 blue 3.5 (Zeiss). After a calibration of the Airyscan detector images are captured with a pixel dwell time of 1-5 µsec, depending on the signal-to-noise ratio within the sample. Airyscan imaging data was further processed using the integrated *3D Airyscan Processing* function within ZEN 3 blue. Further processing was performed by Fiji (ImageJ, version 2.1.0/1.53c).

### Antibody modification

For the conjugation of JF_635_b-succinimidyl ester with amines in the antibody, a mixture of antibody and dye was prepared. First, the desiccated dye was dissolved, using anhydrous dimethylsulfoxide (DMSO, Invitrogen, D12345). Then a 100 µl mixture of 100 µg of the anti-rabbit IgG and a 20-fold molar excess of JF_635_b-succinimidyl ester were incubated together in 200 mM solution of NaHCO_3_ in PBS (1×) (pH 8.1) for 2 h at room temperature. After the incubation, unconjugated dyes were removed and antibodies were transferred into 0.01% NaN_3_ in PBS or just PBS by using Zeba™ Spin Desalting Columns (40K MWCO, ThermoFisher, A57760) following the manufacturer’s suggested protocol.

## Data and Materials availability

All data that support the findings described in this study are available within the manuscript and the related supplementary information, and from the corresponding authors upon reasonable request.

---

**Extended Data Fig. 1.**
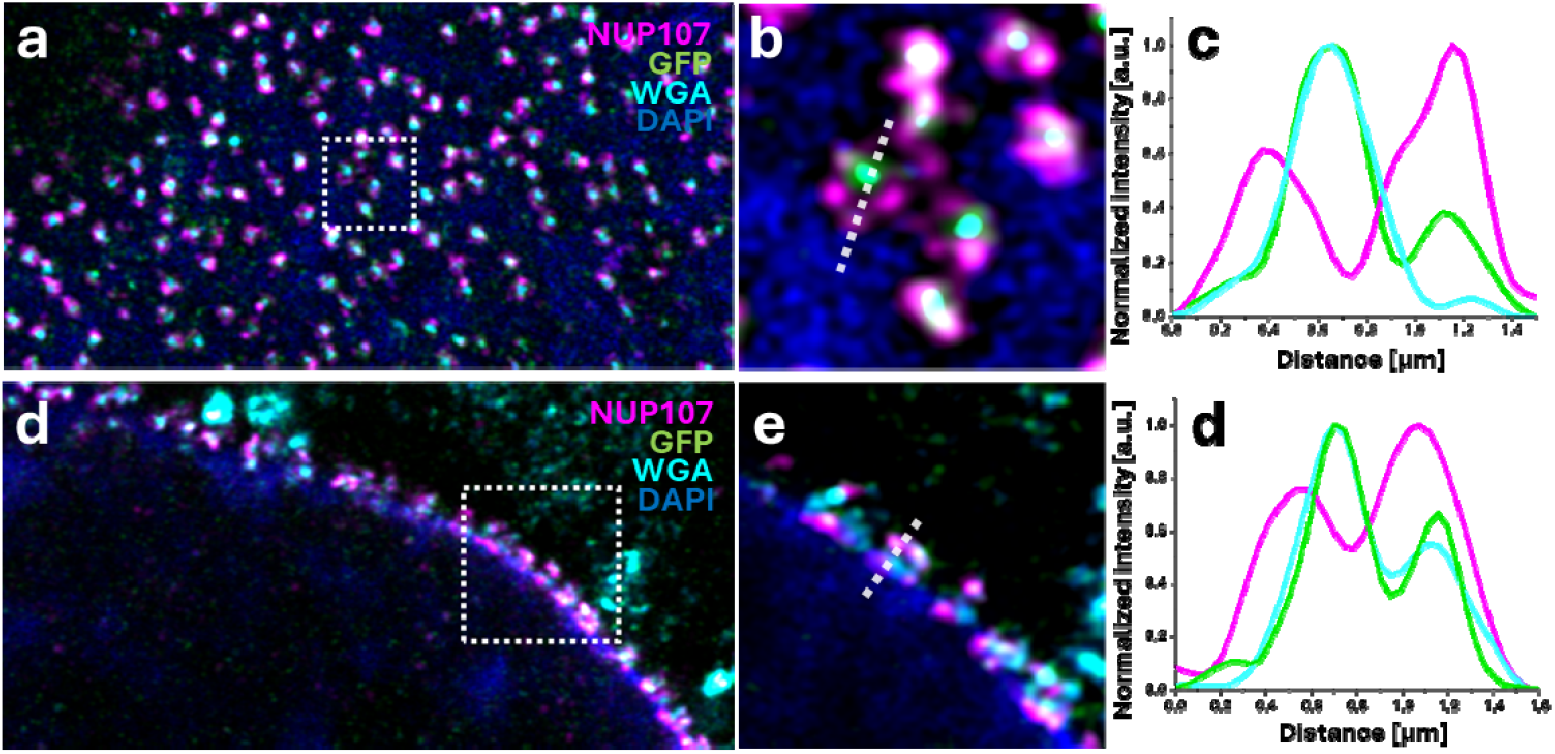
Optimization of digestion enables multicolor imaging with high spatial resolution. **a**, Airyscan image of expanded NPCs of a HeLa-NUP107-GFP cells post-expansion stained with anti-GFP nanobody-X4-ATTO643 (magenta), WGA-CF568 (light blue), NUP107-GFP (green) and DNA with DAPI (blue). **b**, Expanded view of a selected region in (**a**) with the DAPI signal. **c**, Fluorescence intensity plot for GFP and WGA-CF568 disclosing their overlapping signals. **d**, Airyscan image of the equatorial plane of the nuclear lamina, showing a side-profile of expanded NPCs of a HeLa-NUP107-GFP cell stained by nanobody-anti-GFP-X4-ATTO643 (magenta), WGA-CF568 (light blue), NUP107-GFP (green), and DAPI (blue). **e**, Zoom-in of the outlined region in (**d**) showing side-profiles of NPCs. **d**, Cross-sectional profile (yellow dotted line in (**f**)) showing the intensities of the anti-GFP nanobody-X4-ATTO643, GFP, and WGA-CF568. Scale bars: 5 µm (**a**,**d**), 500 nm (**b**,**e**), and 1 µm (**f**).

**Extended Data Fig. 2.**
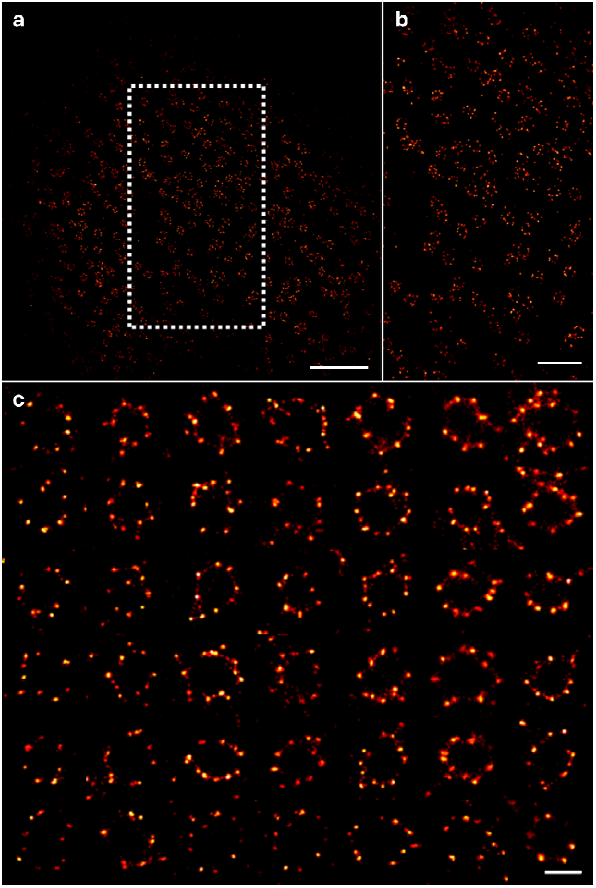
JF_635_b enables SMLM of the NPC in fully expanded hydrogels. **a**, SMLM image of a fully expanded HeLa-Nup107-GFP cell, post-expansion labeled with primary rabbit-anti-GFP antibody and secondary anti-rabbit antibody labeled with JF_635_b. **b**, Expanded view of the outlined region in (**a**). **c**, SMLM images of individual NPCs used for calculation of an average NPC diameter of 659.8 ± 79.6 nm (mean ± SD, n=43) corresponding to an average expansion factor of ∼6.2×. Scale bars: 5 µm (**a**), 3 µm (**b**), and 500 nm (**c**).

**Extended Data Fig. 3.**
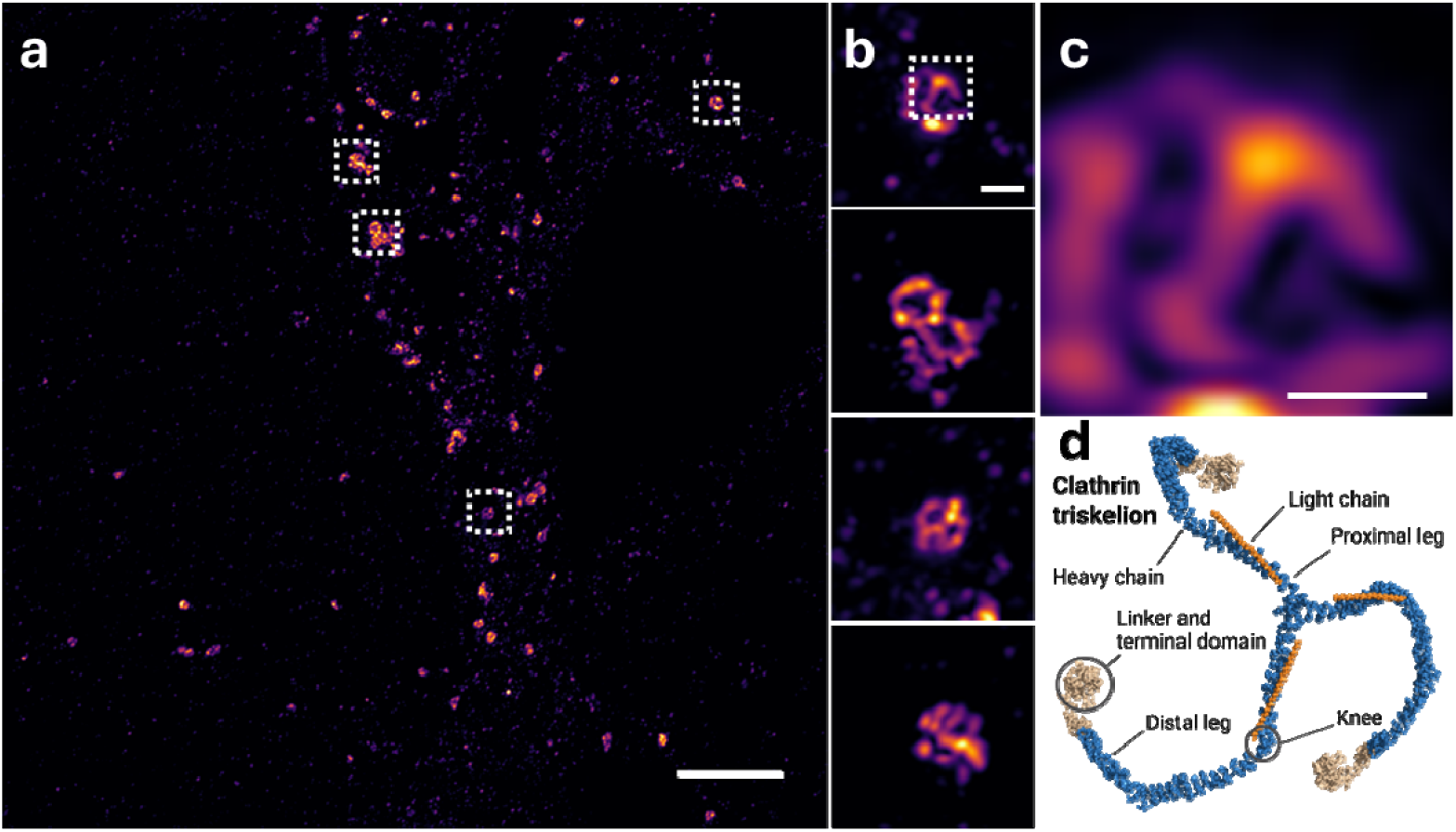
Post-expansion labeled clathrin-coated pits reveal triskelia. **a-c**, Airyscan images of expanded clathrin coated pits stained with conventional primary (pAB) and fluorescently (CF568) labeled secondary IgG antibodies (sAB). **a**, Overview image of multiple clathrin-coated pits in a cell. **b**, Expanded views of individual clathrin-coated pits of the outlined regions in (**a**). **c**, Zoom-in of (**b**) showing a clathrin triskelion. **d**, Schematic representation of the clathrin triskelion. The heavy chain is displayed in blue, the light chain in orange and the terminal domain in beige (PDB 1XI4)^37^. Scale bars: 20 mm (**a**), 1 µm (**b**), and 500 nm (**c**).

