## Supplementary Figures for "Nanoscale imaging of expanded cells and proteins with spontaneously blinking dyes"

### Single-step expansion SMLM enables molecular-resolution imaging in cells and isolated proteins

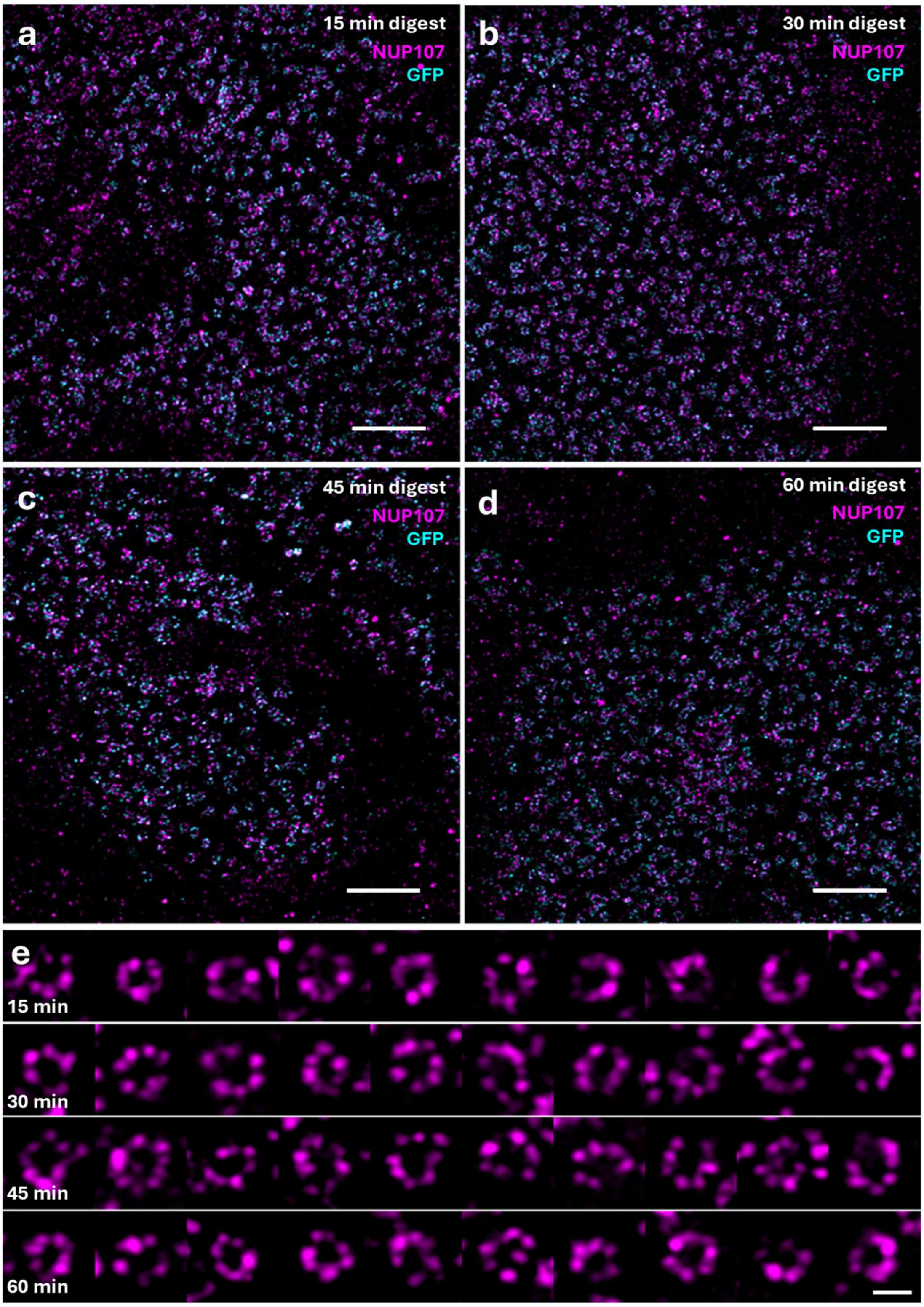


**Supplementary Figure 1. Shortened digestion times preserve NPC structure. a-d,** Airyscan images of expanded HeLa-NUP107-GFP cells in TREx hydrogels, digested with 8 U/ml Proteinase K in optimized 10x digestion buffer for 15 min (**a**), 30 min (**b**), 45 min (**c**), and 60 min (**d**) at 50°C. NPCs were pre-expansion labeled with biotinylated anti-GFP-nanobody and ATTO-643-streptavidin. **e,** Gallery of NPC images from (**a-d**) which were used to measure the diameter via cross sectional profiles. Peak-to-peak distances were measured separately: 0.99 ± 0.10 µm (mean ± s.d.) (n = 12) for 15 min (**a**), 1.04 ± 0.14 µm (mean ± s.d.) ) (n = 15) for 30 min (**b**), 1.08 ± 0.13 µm (mean ± s.d.) ) (n = 12) for 45 min (**c**) and 1.07 ± 0.09 µm (mean ± s.d.) ) (n = 8) for 60 min (**d**). Scale bars: 10 µm (**a-d**), 500 nm (**f**).

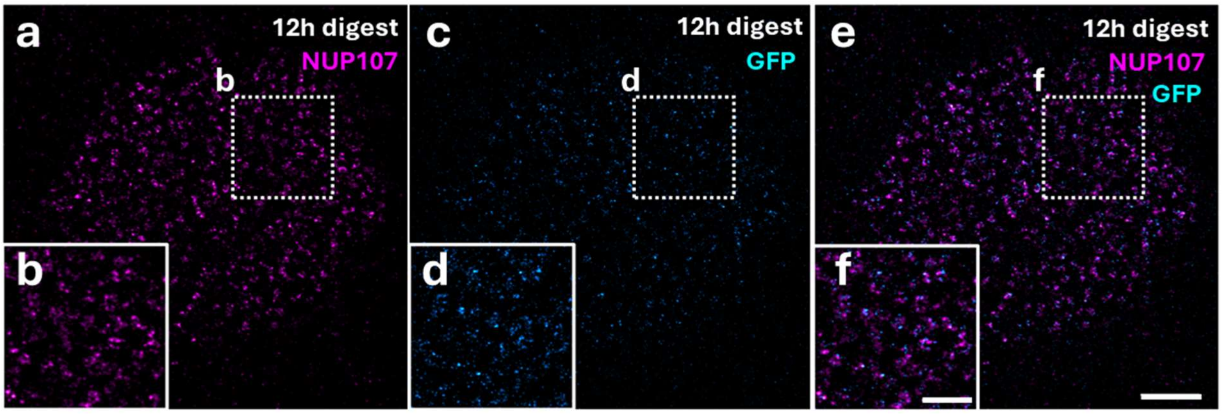


**Supplementary Figure 2. Prolonged digestion times disrupt the structural integrity of NPCs.** Airyscan images of expanded HeLa-NUP107-GFP cells post-expansion labeled by biotinylated anti-GFP-nanobody and ATTO643-streptavidin in TREx hydrogel. Samples were digested with 8 U/ml Proteinase K in 10x digestion buffer for 12 h. **a**,**b,** ATTO643-streptavidin signal. **c, d,** GFP signal. **e, f',** merged image of both signals. Scale bars: 10 µm (**a, c, e**), 5 µm (**b, d, f**).

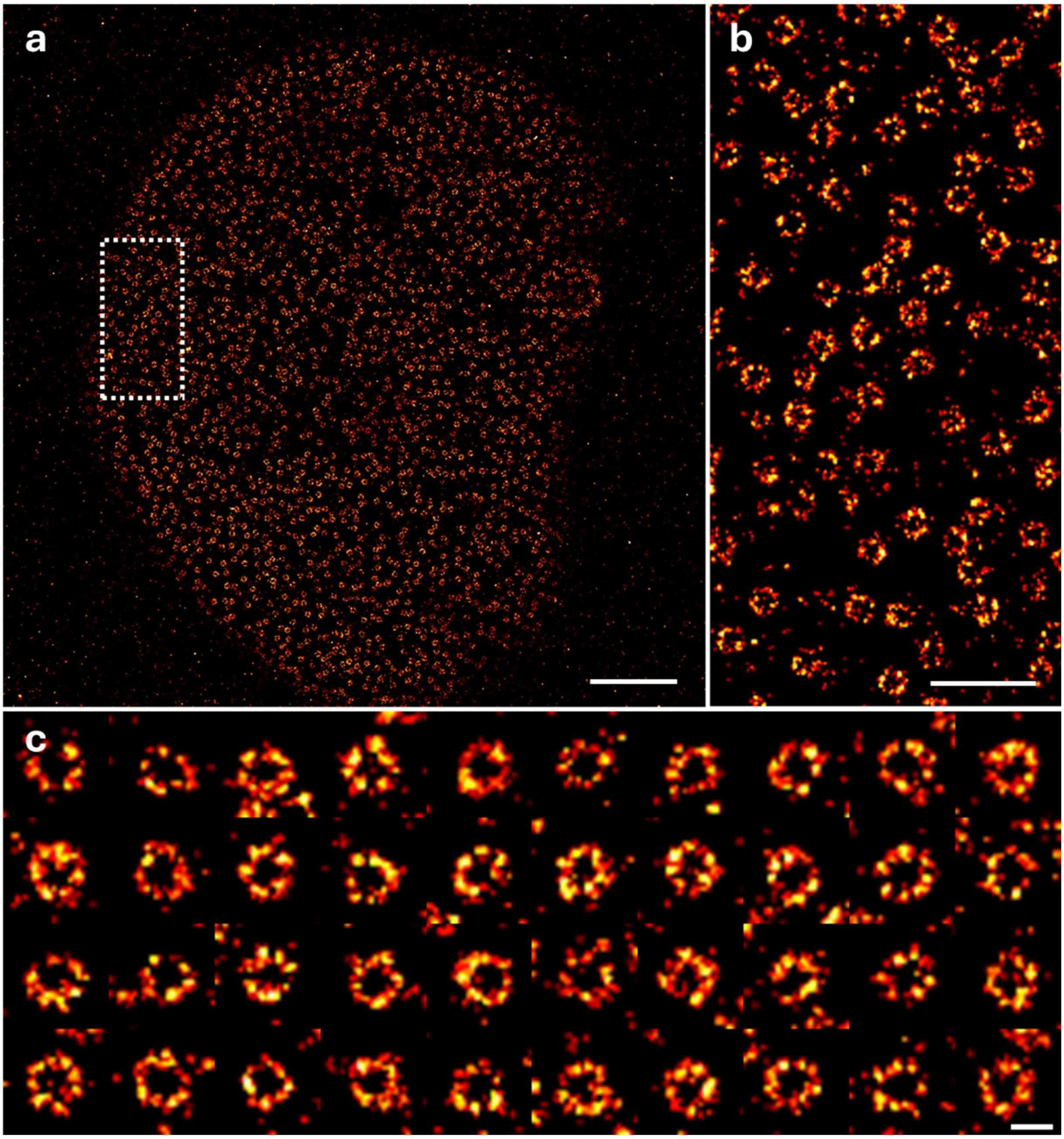


**Supplementary Figure 3. *d*STORM images of unexpanded NPCs. a,** *d*STORM image of the nucleus of a HeLa-NUP107-GFP cell labeled with anti-GFP-nanobody-X4-AF647. **b**, Expanded view of the region marked with a white rectangle in (**a**). **c,** *d*STORM images of individual NPCs used for calculation of an average NPC diameter of 105.8 nm ± 6.8 nm (mean ± s.d., n=42) matching the expected value of 107 nm[1,2]. Scale bars: 3 µm (**a**), 1 µm (**b**), and 100 nm (**c**).

.

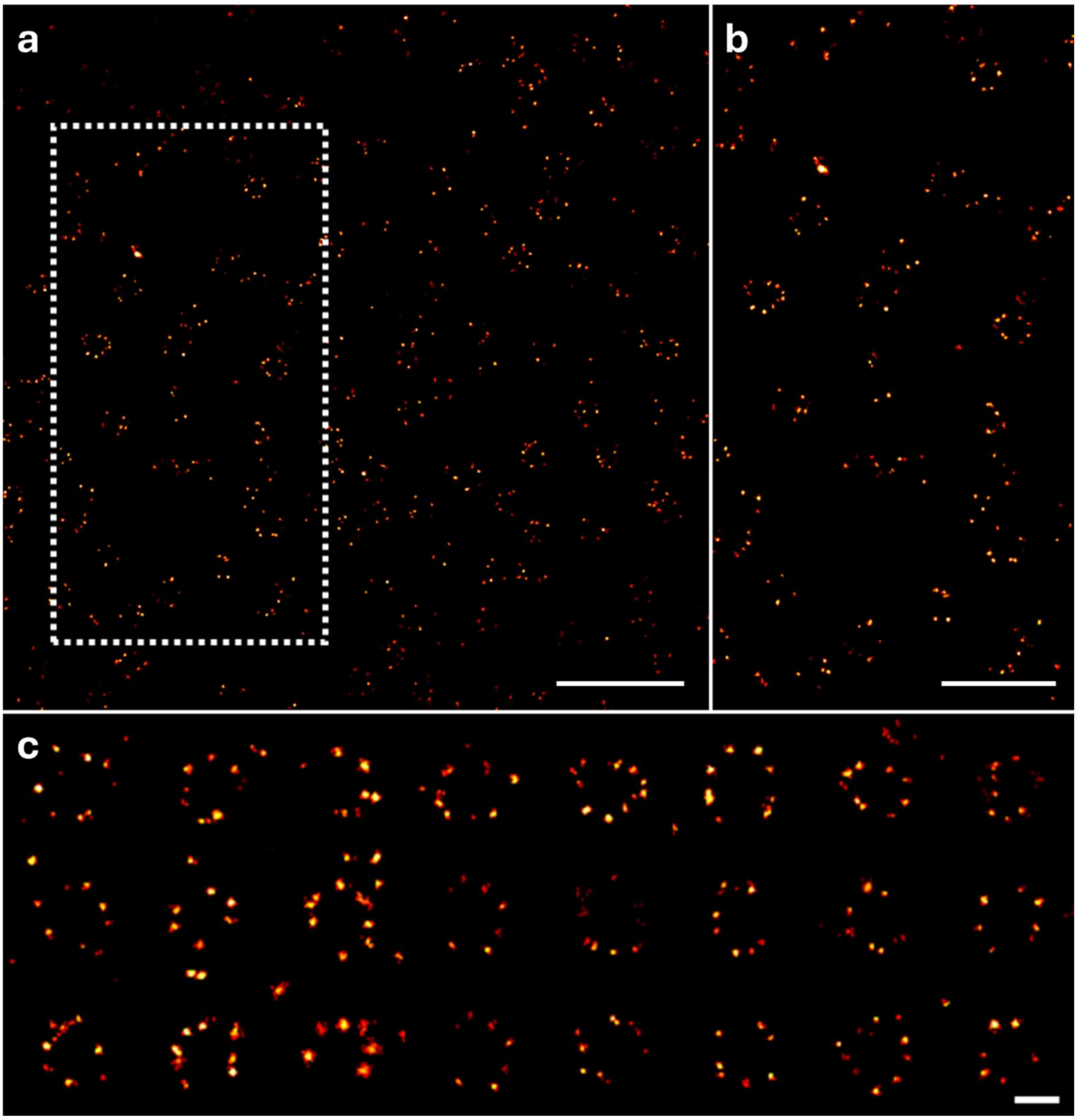


**Supplementary Figure 4. Re-embedding enables Ex-*d*STORM. a,** Ex-*d*STORM image of a HeLa-NUP107-GFP cell post-expansion labeled with anti-GFP-nanobody-X4-AF647. **b,** Zoom-in of the region marked white in (**a**). **c,** Selected NPCs to determine an average peak-to-peak distance of AF47 signals of 564.1 nm ± 50.9 nm (mean ± s.d., n=24) corresponding to an expansion factor of 5.3×. Scale bars: 3 μm (**a**), 2 μm (**b,c**), and 400 nm (d).

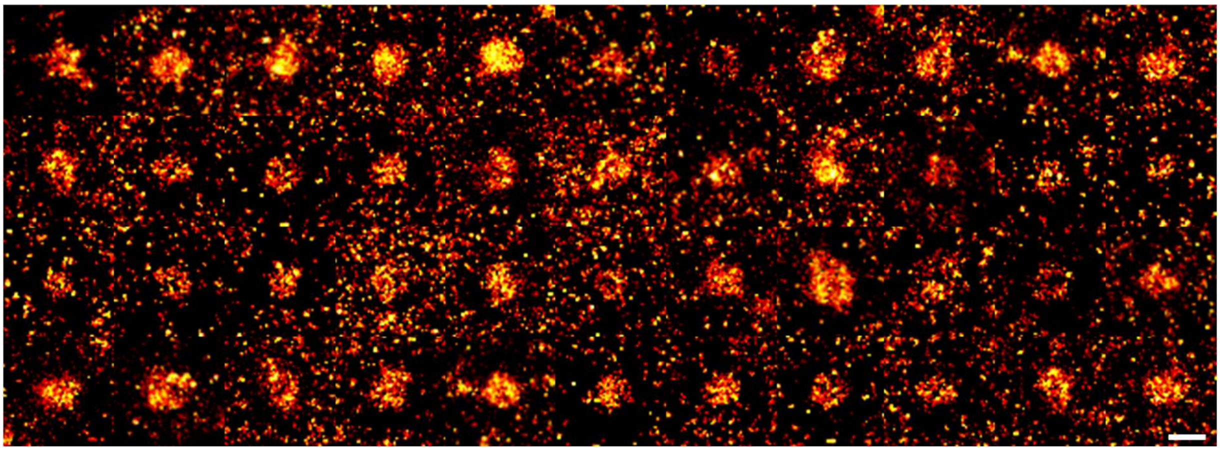

**Supplementary Figure 5. Super-resolution imaging of isolated $JF_{635}$b labeled PCNA.** A gallery of isolated PCNA proteins labeled with $JF6_{35}$b-NHS ester showing single, cropped out PCNA molecules that were averaged. Scale bar: 200 nm.

**Supplementary Movies**

**Supplementary Movie S1.** Blinking movie of an exemplary 256 × 256 pixels area of Ex-SMLM imaging of a fully expanded HeLa-Nup107-GFP cell, post-expansion labeled with primary rabbit-anti-GFP antibody and secondary anti-rabbit antibody with $JF_{635}b$. Movie played at 50 fps.

**Supplementary Movie S2.** Blinking movie of an exemplary 256 × 256 pixels area of single-step Ex-SMLM imaging of a fully expanded hydrogel containing isolated PCNA molecules labeled post-expansion with $JF_{635}b$-NHS. Movie played at 20 fps.
